# Effects of ketamine on GABAergic and glutamatergic activity in the mPFC: biphasic recruitment of GABA function in antidepressant-like responses

**DOI:** 10.1101/2024.07.29.605610

**Authors:** Manoela V. Fogaça, Fernanda Daher, Marina R. Picciotto

## Abstract

Major depressive disorder (MDD) is associated with disruptions in glutamatergic and GABAergic activity in the medial prefrontal cortex (mPFC), leading to altered synaptic formation and function. Low doses of ketamine rapidly rescue these deficits, inducing fast and sustained antidepressant effects. While it is suggested that ketamine produces a rapid glutamatergic enhancement in the mPFC, the temporal dynamics and the involvement of GABA interneurons in its sustained effects remain unclear. Using simultaneous photometry recordings of calcium activity in mPFC pyramidal and GABA neurons, as well as chemogenetic approaches in *Gad1-Cre* mice, we explored the hypothesis that initial effects of ketamine on glutamate signaling trigger subsequent enhancement of GABAergic responses, contributing to its sustained antidepressant responses. Calcium recordings revealed a biphasic effect of ketamine on activity of mPFC GABA neurons, characterized by an initial transient decrease (phase 1, <30 min) followed by an increase (phase 2, >60 min), in parallel with a transient increase in excitation/inhibition levels (10 min) and lasting enhancement of glutamatergic activity (30-120 min). Previous administration of ketamine enhanced GABA neuron activity during the sucrose splash test (SUST) and novelty suppressed feeding test (NSFT), 24 h and 72 h post-treatment, respectively. Chemogenetic inhibition of GABA interneurons during the surge of GABAergic activity (phase 2), or immediately before the SUST or NSFT, occluded ketamine’s behavioral actions. These results indicate that time-dependent modulation of GABAergic activity is required for the sustained antidepressant-like responses induced by ketamine, suggesting that approaches to enhance GABAergic plasticity and function are promising therapeutic targets for antidepressant development.

## Introduction

Major depressive disorder (MDD) is a recurring neuropsychiatric illness that has a lifetime prevalence of ∼17% in the U.S., and is a leading cause of disability worldwide (1). Despite the significant economic and human impact, the effectiveness of treatments remains suboptimal, underscoring MDD’s inherent heterogeneity and our limited comprehension of the molecular and functional mechanisms that underlie its etiology (2). Traditional antidepressants take weeks to months to induce a therapeutic response, and up to 33% of patients prescribed these medications are considered treatment resistant (3). Conversely, low doses of ketamine, an NMDA receptor (NMDA-R) blocker, can induce rapid (2 h) and sustained (up to 7 days) antidepressant effects in patients diagnosed with MDD, even in patients that are refractory to current antidepressant medications (4).

Human and rodent studies indicate that depression and chronic stress are linked to structural alterations in limbic brain areas, including the medial prefrontal cortex (mPFC), characterized by reduced volume, neuronal atrophy, and impaired excitatory synapse density and function (5–9). Conversely, accumulating evidence suggests that ketamine can reverse these deficits and produce rapid antidepressant effects through an initial, transient blockade of NMDA receptors (NMDA-R) in GABA interneurons, leading fast disinhibition of excitatory pyramidal neurons and triggering neuronal plasticity in the mPFC (10–12). An alternative hypothesis is that ketamine acts directly on pyramidal neurons to block NMDA-R activity driven by spontaneous glutamate release (13).

Furthermore, in addition to disruption in excitatory synapses, MDD subjects and animals exposed to chronic stress have reductions in cortical and plasma GABA levels, as well as several GABA markers in the PFC (10, 14–20). Since inhibitory inputs control network excitability, integration, and synchrony, deficits in GABA function compromise the signal-to-noise properties of glutamatergic neurons and the integrity of circuit-level information transmission from the mPFC to projection areas (5). Consistent with this idea, prefrontal cortical GABA abnormalities are associated with hippocampal structural deficits in MDD subjects (9) and normalization of GABA function is associated with remission of depressive symptoms (18, 21–27). Indeed, following the initial enhancement of glutamate function, ketamine and other rapid antidepressants also increase GABA signaling in the mPFC, potentially contributing to reestablishing the integrity of excitatory and inhibitory signal efficiency and precision in corticolimbic circuits (14–16, 19, 20, 28).

In this study, we hypothesized that ketamine might induce long-lasting changes in the activity of GABA interneurons that could contribute to sustained antidepressant effects. To test this, we imaged activity of mPFC pyramidal and GABA neurons simultaneously following ketamine administration and during behavioral tests relevant to antidepressant efficacy. We then employed chemogenetic approaches to investigate whether activity of GABA interneurons is necessary and/or required for the behavioral actions of ketamine.

## Materials and Methods

### Animals

Male glutamic acid decarboxylase 1 (*Gad1*)*-Cre* transgenic mice and WT littermates (8-12-week-old) on the C57BL/6 background were bred in-house as in previous studies (29, 30). Male mice were initially chosen to allow for comparison of the outcomes observed following ketamine administration with previous literature, predominantly described in male mice. All animals were group-housed with a 12/12h light-dark cycle and food and water *ad libitum*. Following surgery for viral infusion and/or optical fiber placement, animals were single housed for 4 weeks and remained isolated until the end of the experiments. All procedures were conducted in compliance with the National Institute of Health (NIH) guidelines for the care and use of laboratory animals and were approved by the Yale Institutional Animal Care and Use Committee.

### Viral constructs and surgery

Adeno-associated viruses AAV.CamKII.GCaMP6s.WPRE.SV40, pAAV.Syn.Flex.NES-jRCaMP1b.WPRE.SV40, ≥ 1 x 10^13^ vg/ml, and AAV2-hSyn-DIO-hM4D(Gi)-mCherry (hM4DGi), ≥ 7 x 10^12^ vg/ml, were obtained from Addgene (USA). To image activity of pyramidal neurons and GABA interneurons simultaneously, anesthetized *Gad1-Cre* mice (ketamine, 100 mg/kg; and xylazine, 10 mg/kg) received unilateral intra-mPFC infusion of a cocktail (1:1; 0.6 µl, 0.1µl/min) containing two calcium sensors with non-overlapping spectra: CaMKII-driven GCaMP6s (GCaMP6s) and Cre-driven jRCaMP1b (RCaMP) viruses (coordinates from bregma: anterior-posterior: +1.9 mm; medial-lateral: ± 0.4 mm; dorsal-ventral -2.7 mm), along with implantation of a fiber (stainless steel ferrule, 400 µm core, 0.50 NA, 2.5 mm length, ThorLabs, Newton, New Jersey, USA) in the same region. The fiber was implanted 0.2 mm above the injection site and maintained in place with adhesive dental cement (C&B-Metabond, Parkell, NY, USA). Animals received i.p. injections of carprofen (5 mg/kg) immediately after the surgery and daily for the next 2 days. Following surgery, all animals remained single-housed throughout the duration of the protocols. In this study, single housing was used to prevent detachment of the optical fiber from the skull and to serve as a chronic mild stressor to study inhibitory and excitatory neuronal activity. To preserve the integrity of the cannula, including a control group not subjected to isolation, which would enable comparing stress effects, was not feasible for practical reasons. For chemogenetic experiments, animals received bilateral infusion of inhibitory hM4DGi (0.5 µl/side; 0.1µl/min) into the mPFC and underwent the same post-surgery protocol as described, including single housing. Fiber placement and viral efficiency were analyzed using a confocal microscope to obtain Z-stack image sequences (Leica TSE-SPE) (30).

### Drug administration

Ketamine (Sigma-Aldrich, 10 mg/kg, i.p.) was dissolved in saline. Clozapine-N-oxide (CNO, 0.5 or 1 mg/kg i.p., Enzo Life Sciences, Farmingdale, NY, USA) was administered at different time points, as indicated, based on our previous studies indicating lack of behavioral and locomotor effects at these low doses (30); however, to control for any off target effects that have been reported at higher doses (5-10 mg/kg) (31, 32), both WT and *Gad1-Cre* groups received CNO.

### Behavioral studies

Animals were habituated to testing rooms 30 min before each experiment. All behavioral tests were video recorded and conducted between 10 a.m. and 4 p.m. Experiments were scored by an experimenter blind to treatments.

*Sucrose Splash Test (SUST):* A 10% sucrose solution was squirted onto the dorsal coat of the mouse, as has been described (33). Grooming time was measured for 5 min.

*Novelty-Suppressed Feeding Test (NSFT):* Mice were food deprived for 16 h and placed in a dimly lit box (40 *x* 40 *x* 25) with a pellet of food in the center; the latency to feed was measured with a time limit of 10 min, as described (30). Immediately after the test, home cage food intake was measured over a 10 min period as a feeding control.

### Immunofluorescence

*Gad-Cre* mice received a Cre-dependent hM4DGi viral infusion into the mPFC, as described, and SST or PV co-localization with Gad-hM4DGi+ cells was assessed using primary antibodies (mouse anti-SST #sc-74556, 1:200, Santa Cruz; rabbit anti-PV #ab181086, 1:1000, Abcam) and appropriate secondary antibodies (AlexaFluor® 488 goat anti-rabbit or AlexaFluor® 647 goat anti-mouse, 1:1000), as described (20, 30). For quantification, sections containing the mPFC were analyzed using a Keyence BZ-X800 microscope equipped with an optical sectioning module (20X magnification). The total number of Gad-hM4DGi^+^, PV or SST neurons, as well as Gad-hM4DGi^+^ co-localized with PV^+^ or SST^+^ cells, were obtained within each section (3 sections/animal). The results were then averaged across animals (n = 3 mice) and expressed as a percentage of co-localized cells (number of co-localized Gad-hM4DGi^+^ and SST or PV cells/total Gad-hM4DGi^+^, PV or SST cells × 100).

### Fiber photometry

Multichannel fluorescent signals were recorded with a Tucker RZ5P processor (Tucker-Davis Technologies, Alachua, FL, USA) controlled by the Synapse software suite to display excitation (465 nm for GCaMP, green, and 560 nm for RCaMP, red) and reference (405 nm, isobestic control) signals, modulated at 531, 330 and 211 Hz, respectively. The LEDs (Doric, Quebec, Canada) were adjusted to the photodetector (Newport 2151, Irvine, CA, USA) with excitation light intensity of ∼20 µW. Light was passed through a minicube (FMC6AE, Doric) that contained excitation and emission filter sets. Animals were tethered to the system via fiber optic patch cord (400 μM core, 0.50 NA, ThorLabs) connected to a head mounted fiber optic cannula via a ceramic sleeve.

### Data processing and analysis

Signals were low pass filtered with a frequency cutoff of 5 Hz using MATLAB. The excitation signal for each channel was analyzed using an adaptive iteratively reweighted Penalized Least Squares (airPLS) algorithm to correct bleaching and motion artifacts, and regressed against the reference control signal, using the protocol developed by Martianova et al., 2019 (34) (available at https://github.com/katemartian/Photometry_data_processing). The change in fluorescence (dF/F) was calculated as dF/F (465 nm or 560 nm signal–fitted 405-nm signal)/fitted 405-nm signal. To standardize signals across animals prior to analysis, results were normalized by z-scoring and expressed as average z-scored dF/F. As it is currently not possible to use an isobestic point for red-shifted sensors (RCaMP), the 405 nm signal was used as a reference for correcting movement artifacts, as described (34). However, due to the nature of photobleaching associated with long-term recordings shown in Figure 1 (∼150 min), RCaMP fluorescence intensity displayed a gradually decreasing trend evident in the vehicle-treated group (Supplementary Figure 1). Therefore, bleaching in this experiment was additionally corrected applying the Matlab linear function *detrend* (y = detrend(x, n), where n = 1; x – y = linear trend) to the vehicle-treated group, and the resultant curve was subtracted from saline and ketamine signals, as described (35). The *detrend* algorithm computes the least-squares fit of a straight line (or composite line for piecewise linear trends) to the data and subtracts the resulting function from the data. For this correction, we assumed that saline had no significant effects on calcium fluorescence over time. Detrending was not required for the shorter recordings described in Figures 2 and 3 (up to 60 min). To evaluate excitation/inhibition levels, baseline or 10-min binned data were transformed into positive values by adding a constant, and the ratio of GCaMP to RCaMP transients (dF/F) was calculated within each animal.

**Figure 1.**
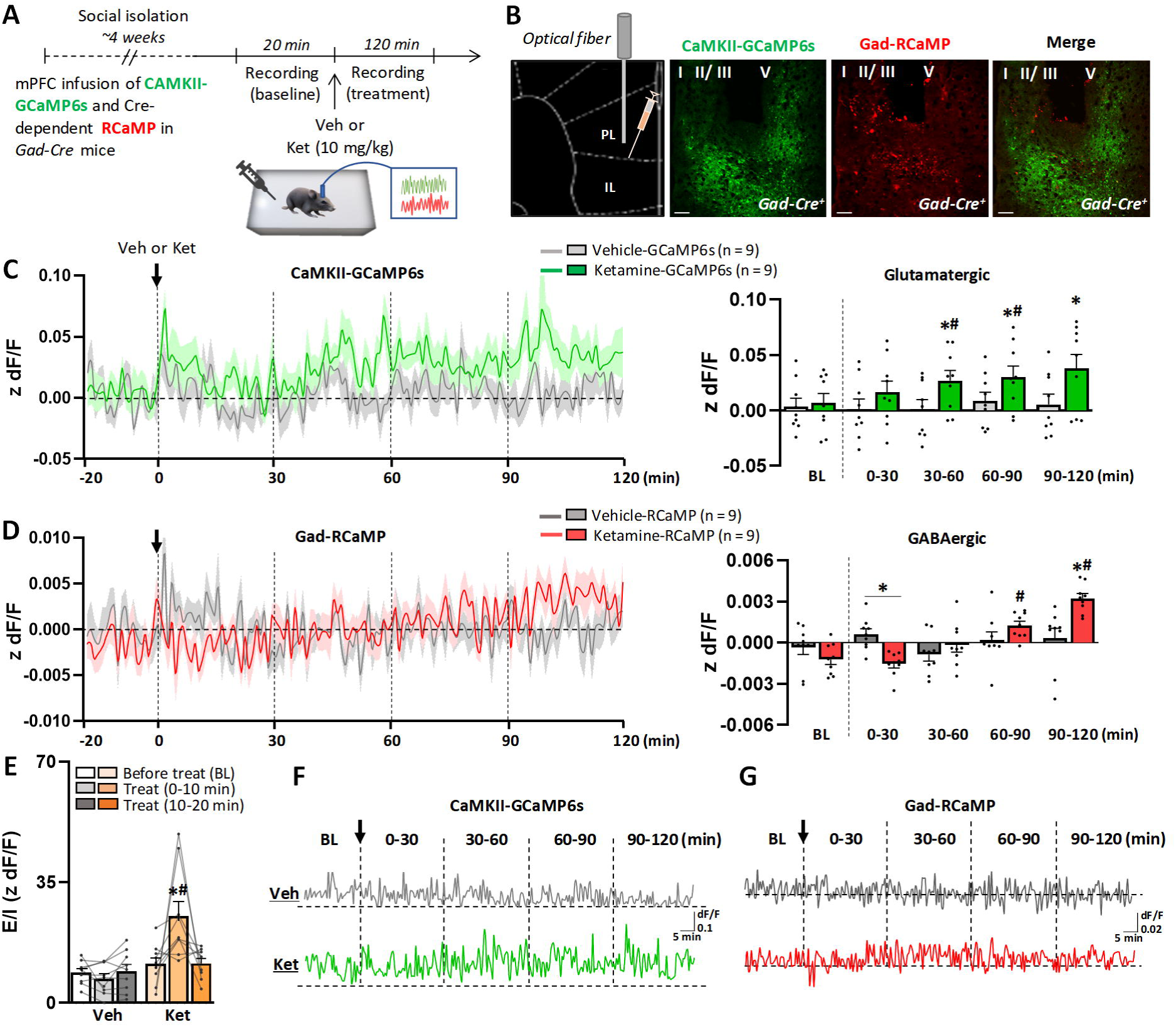
Photometry recordings from mPFC glutamatergic and GABAergic neurons immediately after ketamine treatment. (A) Time course for surgery, treatment and recording. (B) Representative images of the mPFC from *Gad1-Cre* mice showing unilateral infusion of CaMKII-driven GCaMP6s (CaMKII-GCaMP6s) and Cre-dependent jRCaMP1b (Gad-RCaMP) virus, labeling pyramidal and GABAergic cells, respectively, along with implantation of an optical fiber (magnification: 10X). (C) Ketamine increased calcium transients in glutamatergic neurons (CaMKII+) compared to both the baseline (BL, - 20 to 0 min) and vehicle-treated animals 30 min post-treatment, persisting until the end of the recording session (F_treatment 1,8_ = 22.03; F_interaction 4,32_ = 4.99, p ≤ 0.05). (D) Ketamine produced a biphasic response in GABA neuron calcium activity, in which there was an initial decrease (0 to 30 min) followed by an increase in the photometry signal (60 to 120 min) (F_time 4,32_ = 5.23; F_interaction 4,32_ = 9.65, p ≤ 0.05). (E) Ketamine increased excitation/inhibition (E/I) levels within 10 min of administration, which returned to baseline levels shortly thereafter (F_treatment 1,8_ = 27.43; F_interaction 2,16_ = 6.22, p ≤ 0.05). (F) Minimally processed traces representing pyramidal neurons (CaMKII+) expressing GCaMP6s and (G) GABA neurons (Gad-Cre) expressing RCaMP in the mPFC from a vehicle- or ketamine-treated *Gad1-Cre* mouse. The black arrow indicates the treatment time. Horizontal dashed lines indicate z dF/F = 0. Photometry results per animal are computed as average z-scored dF/F, and graphs are presented as mean ± standard error. Repeated measures ANOVA followed by Sidak correction for multiple analysis; *p ≤ 0.05 in comparison to the vehicle-treated group; #p ≤ 0.05 in comparison to the baseline (before treatment), n = 10/group. F_interaction_ represents the interaction between time and treatment factors.

**Figure 2.**
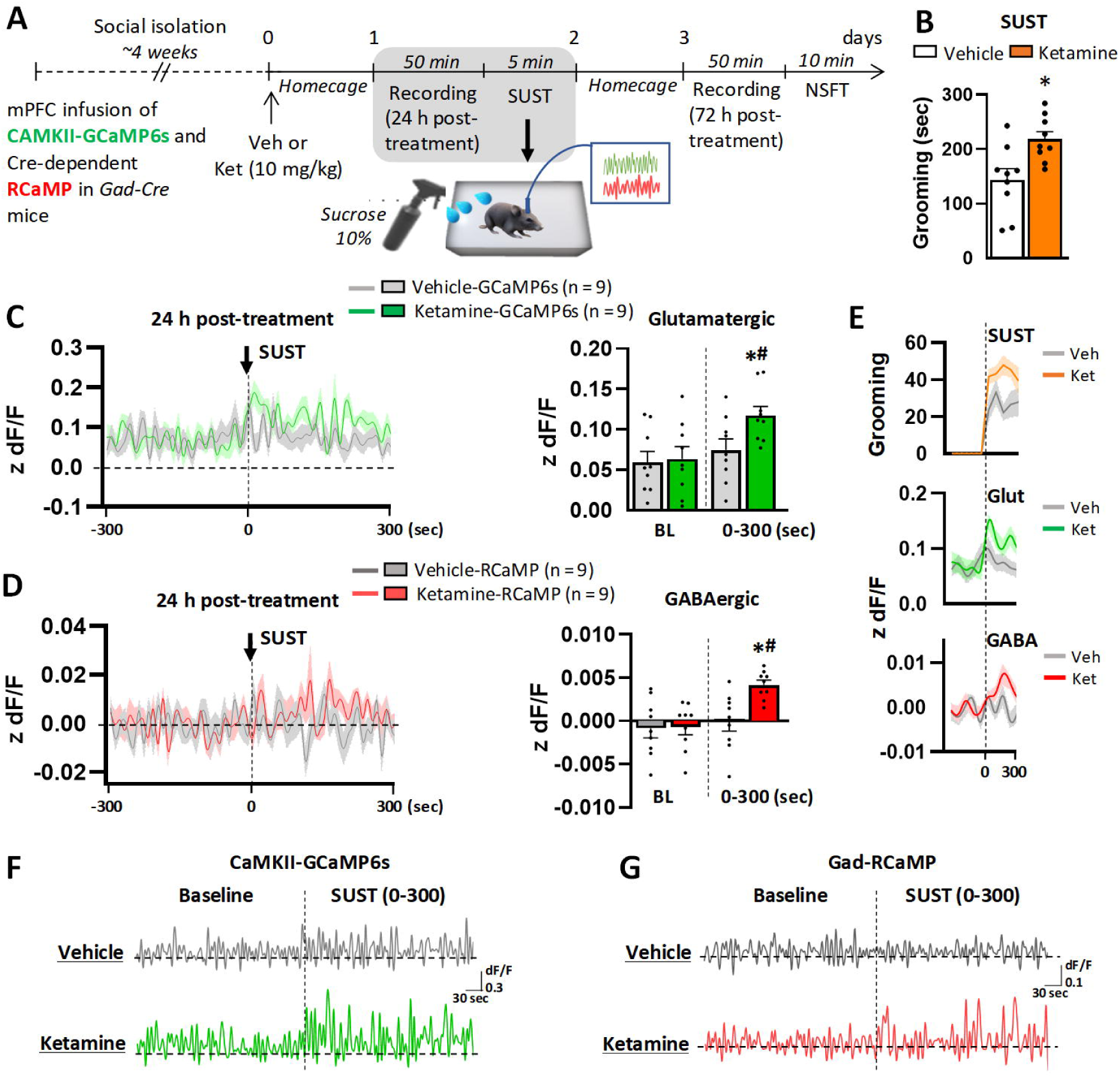
Photometry recordings from glutamatergic and GABAergic neurons 24 h after ketamine treatment and during the sucrose splash test. (A) Timeline for surgery, treatment, recording and behavioral testing. (B) Ketamine increased the grooming time in the sucrose splash test (SUST) 24 h after administration (t_16_ = 3.01, p ≤ 0.05). (C-E) No change in baseline (BL, -300 to 0 sec) activity of pyramidal or GABA neurons was observed 24 h after ketamine administration (p > 0.05). (C) There was an enhancement of calcium transients in both pyramidal (F_time 1,8_ = 25.16; F_interaction 1,8_ = 6.51, p ≤ 0.05) and (D) GABA neurons (F_time 1,8_ = 9.63; F_interaction 1,8_ = 24.86, p ≤ 0.05) in ketamine-treated animals during the SUST (0 to 300 sec) compared to both the baseline and vehicle groups. (E) 60-sec binned representation of the grooming time (orange), GCaMP6s (green, z dF/F) and RCaMP (red, z dF/F) during the SUST duration. (F) Representative traces of pyramidal neurons (CaMKII+) expressing GCaMP6s and (G) GABA neurons (Gad-Cre) expressing RCaMP in the mPFC from vehicle- and ketamine-treated *Gad1-Cre* mice before and during the SUST. Photometry results per animal are computed as average z-scored dF/F, and graphs are presented as mean ± standard error. Repeated measures ANOVA followed by Sidak correction for multiple analysis; *p ≤ 0.05 in comparison to the vehicle-treated group; #p ≤ 0.05 in comparison to the baseline (before test), n = 9/group. F_interaction_ represents the interaction between time and treatment factors.

**Figure 3.**
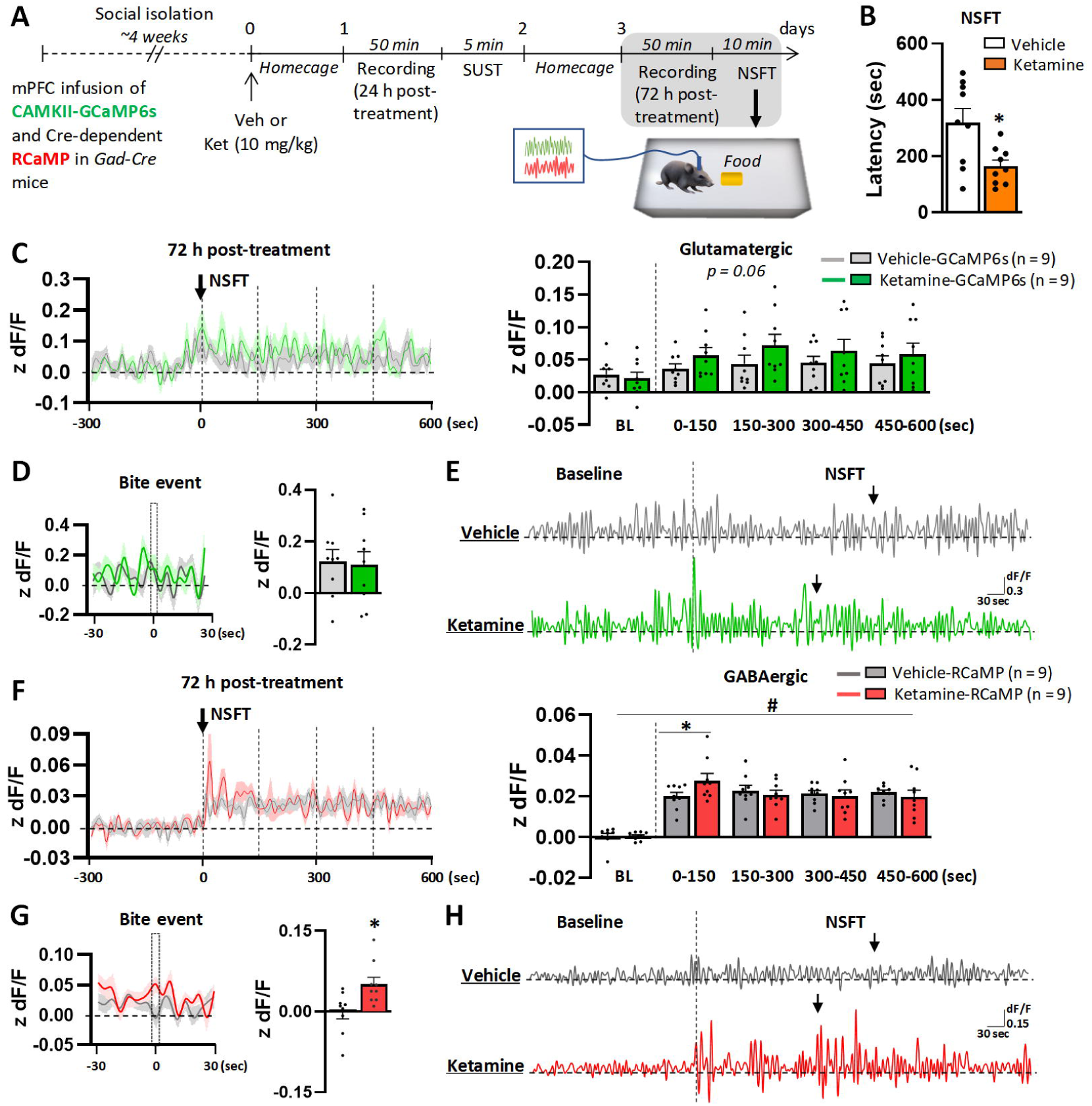
Photometry recordings from glutamatergic and GABAergic neurons 72 h after ketamine treatment and during the novelty suppressed feeding test. (A) Timeline for surgery, treatment, recording and behavioral testing. (B) Ketamine decreased the latency to feed in the novelty suppressed feeding test (NSFT) 72 h after administration (U = 16, p ≤ 0.05). (C) Ketamine did not change baseline (BL, -300 to 0 sec) activity of pyramidal cells 72 h after administration (p > 0.05) and produced a trend towards increased pyramidal activity during the NSFT (150-sec time blocks; vehicle: χ^2^ = 2.22, p > 0.05; ketamine: χ^2^ = 9.24, p = 0.06). (D) No change was observed during the bite event (t_16_ = 0.19, p > 0.05). (E) Representative traces of pyramidal neurons (CaMKII+) expressing GCaMP6s in the mPFC from vehicle- and ketamine-treated *Gad1-Cre* mice before and during the NSFT. (F) Ketamine did not change baseline activity of GABA neurons 72 h after administration (-300 to 0 sec, p > 0.05). However, exposure to the NSFT resulted in an enhancement of GABA neuron activity in both vehicle- and ketamine-treated groups (F_time 4,32_ = 58.83; F_interaction 4,32_ = 3.23, p ≤ 0.05), with a more robust activity observed in the ketamine-compared to the vehicle-group in the first 150 sec of the test (0-150 sec, p ≤ 0.05). (G). Ketamine increased calcium activity in GABA neurons during the bite event (t_16_ = 2.45, p ≤ 0.05). (H) Representative traces of GABA neurons (Gad-Cre) expressing RCaMP in the mPFC from vehicle- and ketamine-treated *Gad1-Cre* mice before and during the NSFT. Black arrows represent a bite event. Photometry results per animal are computed as average z-scored dF/F and graphs are presented as mean ± standard error. Repeated measures ANOVA followed by Sidak correction for multiple analysis, Friedman’s two way analysis of variance by ranks or Mann-Whitney U test; *p ≤ 0.05 in comparison to the vehicle-treated group; #p ≤ 0.05 in comparison to the baseline (before test), n = 9/group. F_interaction_ represents the interaction between time and treatment factors.

In the behavioral studies, z-scores of each epochs of interest (*e.g.,* time blocks following ketamine treatment and during behavioral tests, and the bite event) were averaged across groups and presented in bar graphs. In the NSFT, to account for variability in the time it takes for different animals to bite the food (e.g., to remove a chunk of food from the pellet), the bite event was defined as a 5-second period encompassing the food interaction and bite.

### Statistical analysis

Results were subjected to repeated measures ANOVA (treatment *x* time) with Sidak’s correction for multiple analyses. Additionally, Student’s two-tailed t-test or Two-way ANOVA (treatment *x* genotype) followed by Duncan test were employed as appropriate. All distributions were tested for homogeneity of variance using Levene’s test and for normality using the Kolmogorov-Smirnov test. Non-normal distributions were analyzed by Mann-Whitney U test (two-tailed) or Friedman’s two-way analysis of variances by ranks test. Sphericity was assessed by Mauchly’s test. In cases where the assumption of sphericity was violated, the data were analyzed using the Greenhouse-Geisser correction. Across all analyses, differences were considered significant at p ≤ 0.05. Sample sizes were chosen based on previous experience with the tests employed and power analyses (Cohen’s d power analysis, > 0.8 effect size) conducted following a pilot study. For all analyses, we used the SPSS Software (v 29.0) or GraphPrism (v 9.5.1). The specific test used for each experiment is described in the figure legend. Each experiment was replicated a minimum of 2 times.

## Results

### Photometry recordings immediately after treatment: ketamine enhances activity of excitatory mPFC neurons and elicits a biphasic response in GABAergic neurons

The rapid antidepressant actions of ketamine are thought to involve initial inhibition of GABA interneurons, leading to a subsequent glutamate burst and enhancement of glutamatergic activity in the mPFC (12, 30). Additionally, ketamine administration increases GABA-related synaptic proteins in the mPFC 24 h post-administration (20). However, the temporal dynamics of excitatory and inhibitory neuronal activity in the mPFC following ketamine administration has not been measured. To address this, we infused CaMKII-driven GCaMP6s (green) and Cre-driven jRCaMP1b (RCaMP, red) viruses into the mPFC of *Gad1-Cre* mice to record calcium activity from both glutamatergic (CaMKII^+^) and GABAergic (Gad^+^) neurons simultaneously (Figure 1A). Although previous studies indicate that the CaMKII promoter can lead to unspecific expression in inhibitory neurons (36, 37), our analysis suggest no significant overlap (Figure 1B). Using this dual-channel strategy, we observed that ketamine administration increases the activity of CaMKII^+^ neurons beginning approximately 30 min after administration, and this response persists until the end of the recording period (120 min, Figure 1C, F). In contrast, ketamine decreases the activity of GABA neurons for the first 30 min after administration compared to the vehicle-treated group. It is noteworthy that, since the injection procedure initially produced some level of GABA activity evident in the vehicle-treated group during the first 10 min post-injection (Supplementary Figure 2B), the reduction in calcium transients produced by ketamine becomes apparent only when compared with the vehicle group but not with the baseline (Figure 1D). This response is inverted after 60 min, such that there is a delayed increase in GABA activity compared to baseline (60 min) and to the baseline and vehicle-treated groups (90-120 min) (Figure 1D, G). Finally, because GABAergic and glutamatergic activity shift in opposite directions immediately after ketamine treatment (Supplementary Figure 2), we conducted further analysis to investigate the initial dynamics of ketamine’s actions on E/I levels within each animal. Given that ketamine reaches peak plasma and brain levels within 10 min of intraperitoneal administration in mice (38), we assessed E/I activity during baseline and within 10 or 20 min post-treatment. Interestingly, our results revealed a rapid and transient increase in E/I transients following ketamine administration (10 min) in comparison to its baseline and the vehicle-treated group, which returned to baseline levels shortly thereafter (Figure 1E). These data show that ketamine triggers a biphasic response in GABA interneurons, characterized by a transient decrease followed by an increase of activity.

### Ketamine enhances activity of mPFC pyramidal and GABAergic neurons 24 h post-treatment during the sucrose splash test

Next, we investigated whether ketamine-induced rapid changes in glutamatergic and GABAergic activity are long-lasting and engaged during stress-relevant behavioral tests. An independent cohort of *Gad1-Cre* mice underwent surgery for virus infusion, along with implantation of an optical fiber, as outlined. Twenty-four hours after vehicle or ketamine treatment, animals underwent recording in the home cage to evaluate sustained effects of ketamine in the absence of behavioral challenge (50 min, Supplementary Figure 3) and then during the SUST (5 min) (Figure 2A). Ketamine administration did not alter baseline activity of GABAergic or glutamatergic neurons 24 h after administration (50-min recording, Supplementary Figure 3A-B). For visualization and statistical analyses, baseline traces in Figure 2 represent the last 300 sec of baseline recordings. Ketamine increased grooming time during the SUST (300 sec) (Figure 2B), and increased calcium transients in both glutamatergic (Figure 2C, E-F) and GABAergic cells (Figure 2D, E, G) in comparison to respective baselines and vehicle-treated groups.

### Ketamine enhances activity of GABAergic neurons 72 h post-treatment during the novelty suppressed feeding test

Animals then underwent recordings 72 h following ketamine treatment to evaluate calcium transients in the home cage (50 min) and during the NSFT (10 min) (Figure 3A). As expected, ketamine decreased the latency to feed in the NSFT 72 h following treatment compared to the vehicle-treated group (Figure 3B). No change was observed in the home cage food consumption (Supplementary Figure 4). The results indicate that ketamine did not alter baseline activity of GABAergic or glutamatergic neurons 72 h after administration (0-50 min) (Supplementary Figure 4A-B). For visualization and statistical analyses, baseline traces in Figure 3 represent the last 300 sec of baseline recordings. During the NSFT, ketamine administration led to a strong trend towards increased calcium activity in glutamatergic cells compared to the baseline condition, although it did not reach statistical significance (p = 0.06, Figure 3C). Conversely, there was an increase in calcium transients in GABAergic neurons during the NSFT in both vehicle- and ketamine-treated groups compared to the respective baseline groups (Figure 3F). However, ketamine produced a more robust increase in GABAergic activity during the first 150 s of test duration compared to the vehicle-treated group (Figure 3F). We also measured the average z-scored dF/F paired specifically to the bite event (Figure 3D and G). Interestingly, these results show that ketamine treatment results in activation of GABA interneurons but not pyramidal neurons during the bite event, suggesting that ketamine-induced plasticity in GABAergic neurons is long-lasting and manifest during an avoidance-related behavioral task.

### Chemogenetic inhibition of mPFC GABA neurons following ketamine treatment or during behavioral tests at 24 h or 72 h occludes ketamine’s behavioral actions

Based on the biphasic GABAergic activity observed after ketamine treatment (Figure 1), characterized by an initial decrease (phase 1) followed by an increase (phase 2) in calcium transients, we sought to investigate whether this delayed enhancement of GABA function is required for the sustained behavioral responses induced by ketamine. To address this, *Gad1-Cre* and WT littermate controls were infused bilaterally with a Cre-dependent hM4DGi virus (Figure 4A-B) to selectively inhibit the activity of GABA interneurons in the mPFC during phase 2. To investigate whether the hM4DGi virus expression is restricted to specific subpopulations of interneurons or encompasses overall GABAergic cells in *Gad-Cre* mice, we conducted immunostaining to co-label Gad-hM4DGi^+^ cells with PV or SST. Our results indicate that 30.7% of the Gad-hM4DGi^+^ cells express SST, while 27.1% express PV, revealing heterogeneity within the *Gad-Cre* line (Figure 4C; Supplementary Figure 5). Although PV is shown to be more expressed than SST in the neocortex (40% *vs* 30%, respectively) (39), most of our injections are confined to layers II/III and, to a lesser extent, V of the mPFC, where PV and SST cells are shown to be similarly expressed (39). Conversely, 18.1% and 25.6% of total labeled PV or SST cells, respectively, were Gad-hM4DGi^+^.

**Figure 4.**
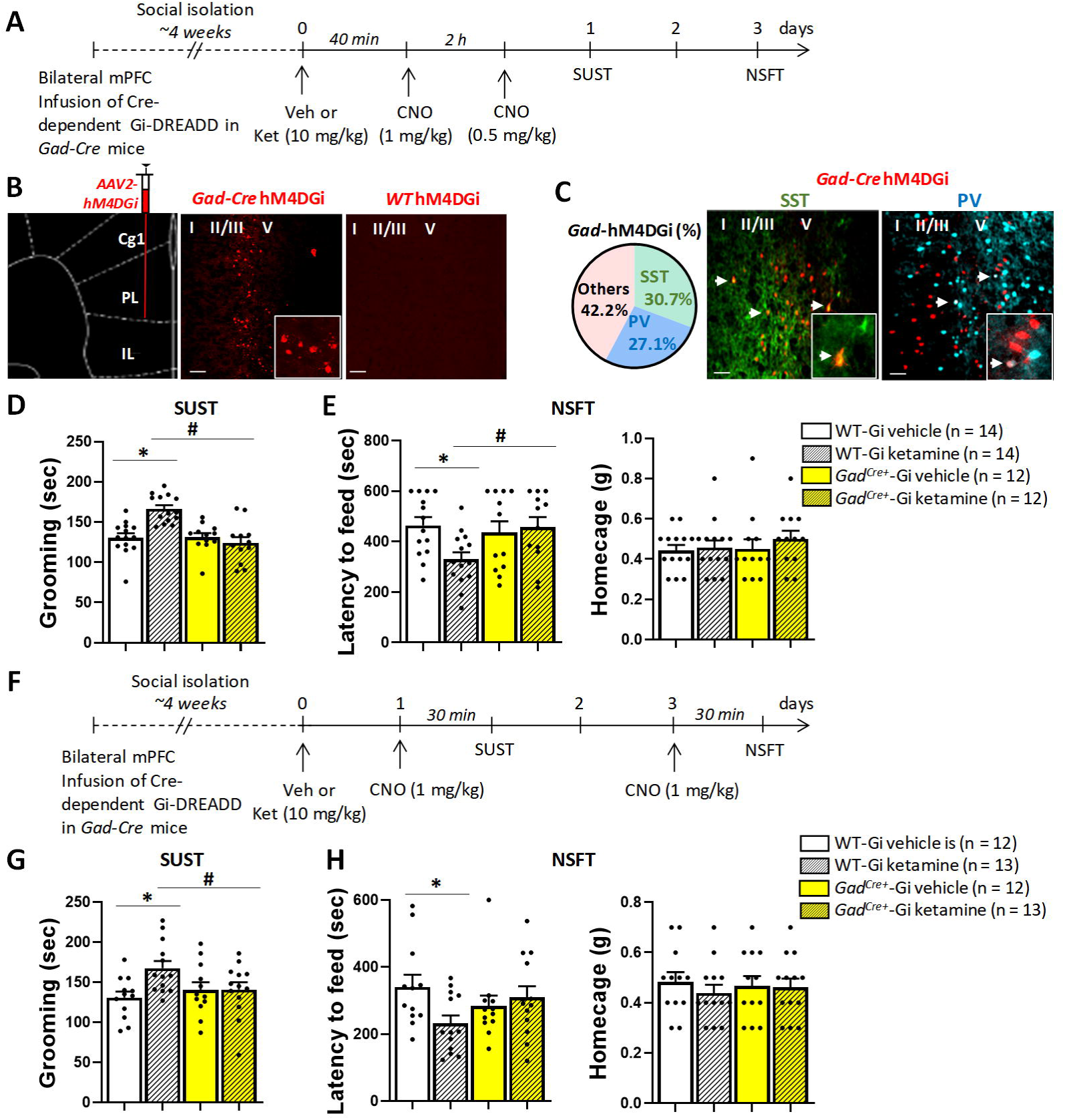
GABA neuron activity is necessary for the sustained behavioral actions of ketamine. (A) Timeline for surgery, treatments, and behavioral testing. (B) Representative images of the mPFC from *Gad1-Cre* and WT (Cre-negative littermate) mice that received bilateral infusions of hM4DGi virus (magnification: 10X; inset: 60X). (C) Co-labeling percentage of PV or SST cells with hM4DGi virus in mPFC sections from *Gad-Cre* mice (magnification: 20X; inset: 40X). (D) CNO administration 40 min (1 mg/kg) and 2 h (0.5 mg/kg) post-ketamine treatment blocked its behavioral effects in the sucrose splash test (SUST, 24 h post-ketamine; F_genotype 1,3_ = 13.42, F_treatment 1,3_ = 6.33, F_interaction 1,3_ = 14.52, p ≤ 0.05) and (E) novelty suppressed feeding test (NSFT, 72 h post-ketamine; F_interaction 1,3_ = 4.67, p ≤ 0.05). There was no change in home cage food consumption (F_interaction 1,3_ = 0.22, p > 0.05). (F) Timeline for surgery, treatments, and behavioral testing. (G) CNO administration (1 mg/kg) 30 min before the behavioral test occluded the behavioral effects of ketamine in the sucrose splash test (SUST, 24 h post-ketamine; F_treatment 1,3_ = 4.34, F_interaction 1,3_ = 4.21, p ≤ 0.05) and (H) novelty suppressed feeding test (NSFT, 72 h post-ketamine; F_interaction 1,3_ = 4.66, p ≤ 0.05). There was no change in home cage food consumption (F_interaction 1,3_ = 0.30, p > 0.05). Graphs are presented as mean ± standard error. Two-way ANOVA followed by Duncan; *p < 0.05 in comparison to the WT vehicle-treated group; #p < 0.05 in comparison to the WT ketamine-treated group. F_interaction_ represents the interaction between genotype and treatment factors.

In this experimental setup, animals received either vehicle or ketamine (10 mg/kg), followed by CNO (1 mg/kg) 40 min later (Figure 4A). This schedule of administration allows the initial ketamine-induced blockade of GABA interneurons to occur (<30 min, phase 1), while occluding the subsequent enhancement of GABA activity (phase 2). CNO has a short half-life in mice (∼1 h) (31), so an additional low dose of CNO (0.5 mg/kg) was administered after 2 h to extend the effect. It is noteworthy that these low doses of CNO do not elicit behavioral or locomotor responses 24 h or 72 h later when injected in hM4DGi-infused *Gad1-Cre* mice (30). Ketamine increased grooming time in the SUST (24 h) and decreased the latency to feed in NSFT (72 h) in WT mice, as expected, and these effects were blocked by chemogenetic inhibition of GABA interneurons during phase 2 in *Gad1-Cre* mice (Figure 4D-E). No changes were observed in home cage food consumption (Figure 4D). This suggests that activity at GABAergic synapses is required for the sustained antidepressant-like effects of ketamine. In a separate control experiment, we adopted an inverse approach where hM4DGi-infused *Gad1-Cre* mice received CNO (1 mg/kg) 40 min *before* vehicle or ketamine, and then were tested in the SUST or NSFT, 24 or 72 h post-treatment, respectively (Supplementary Figure 6). This protocol allows us to evaluate CNO pre-treatment on ketamine’s actions without affecting the delayed GABAergic surge, as CNO would be largely eliminated by this time. As expected, prior CNO treatment did not impact ketamine’s behavioral effects in both the SUST (Supplementary Figure 6C) and NSFT (Supplementary Figure 6D), supporting our hypothesis that ketamine-induced GABAergic activity is time-dependent and engaged at a later timepoint.

Based on the observation that GABAergic transients are increased *during* the SUST and NSFT tests 24 and 72 h following ketamine treatment, respectively (Figure 2 and 3), we determined whether activation of GABA interneurons *during* these tests is required for the behavioral outcomes induced by ketamine. WT and *Gad1-Cre* mice were administered either vehicle or ketamine, followed by CNO (1 mg/kg) 24 and 72 h later, injected 30 min before each behavioral test (Figure 4F). Importantly, CNO does not produce any behavioral or locomotor effects when administered 30 min before behavioral tests in hM4DGi-infused *Gad1-Cre* mice (30). Previous ketamine administration increased grooming time in the SUST (24 h after administration) and decreased the latency to feed in NSFT (72 h after administration) in WT mice, and these effects were occluded by chemogenetic inhibition of GABA interneurons 30 min before each test (Figure 4G-H). No differences were observed in home cage food consumption among groups (Figure 4H). This suggests that ketamine-induced activity of GABAergic neurons is long-lasting and engaged during stress-related behavioral tasks.

## Discussion

In this study we provide direct evidence that ketamine exerts sustained antidepressant-like actions through facilitation of GABA function. The results support the hypothesis that ketamine modulates the activity of mPFC GABA interneurons in a biphasic manner, with an initial decrease in activity (phase 1), accompanied by activation of pyramidal neurons, followed by an increase in GABA function (phase 2) (Figure 5). Notably, chemogenetic data indicate that this delayed enhancement of mPFC GABA activity is both required and necessary for the sustained behavioral actions of ketamine. The increase in GABA signaling after ketamine administration is also long-lasting and engaged during behavioral tests relevant to stress responses, suggesting it may be involved in fine-tuning cortical circuits during stress-related behaviors.

**Figure 5.**
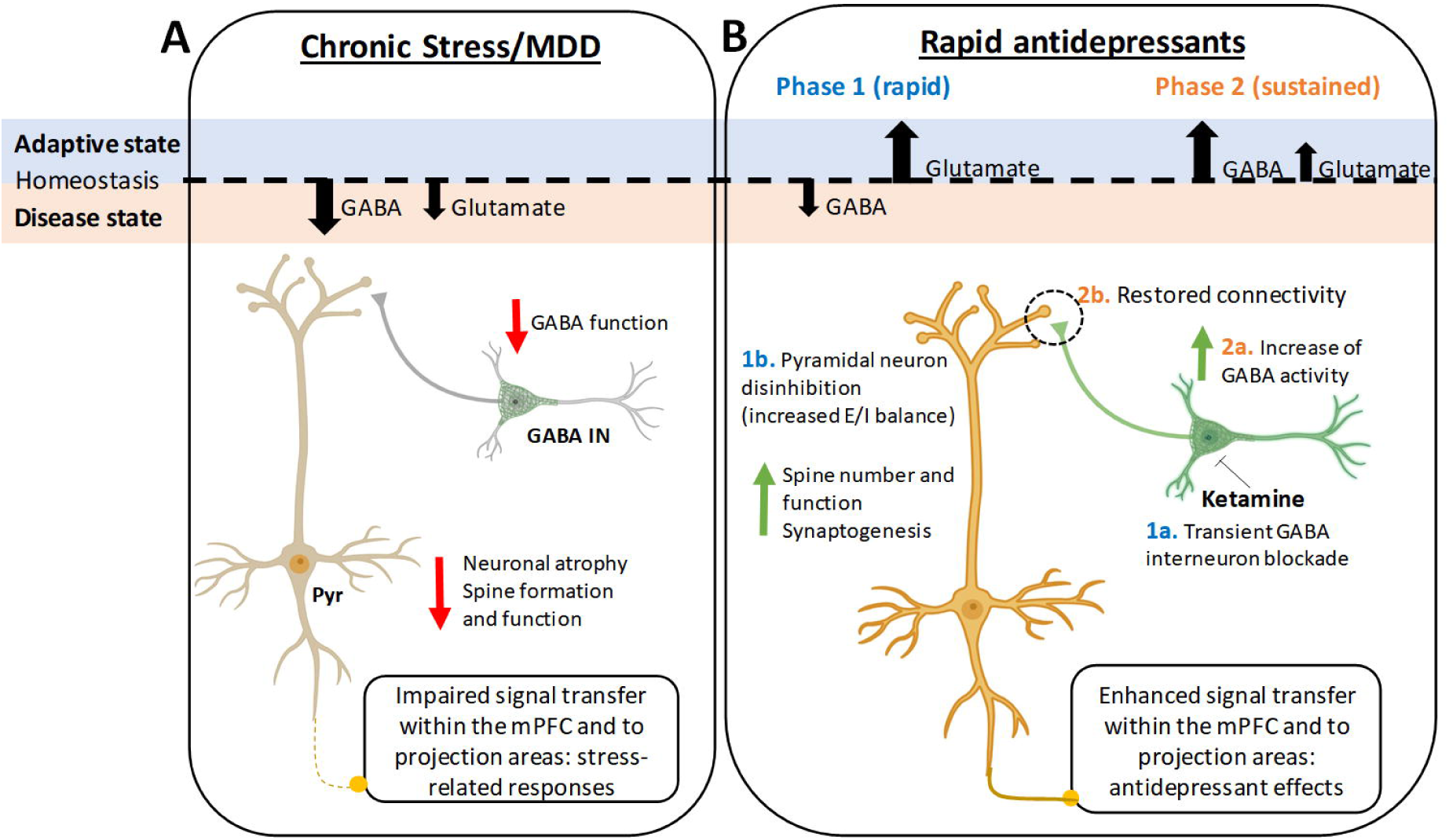
Biphasic glutamatergic-GABAergic model for ketamine action. (A) Major depressive disorder (MDD) and chronic stress impair glutamate and GABA function in the mPFC, leading to altered connectivity and network dysfunction in corticolimbic brain regions, as suggested by previous studies (5, 14, 18, 20, 60, 69). (B) Ketamine restores these deficits by acting through two phases: (1a) In phase 1 (acute), ketamine blocks the activity of GABA interneurons via NMDA-R, disinhibiting pyramidal cells and generating a glutamate burst and increasing excitation/inhibition (E/I) levels (1b). The glutamate burst triggers a cascade of cellular events culminating in synaptic plasticity and fast antidepressant responses, as suggested by previous studies (12, 20, 30, 69). In a second phase (sustained), there is a self-tuning adjustment to reach E/I balance in the mPFC, leading to potentiation of GABAergic function (2a) and restored circuit connectivity (2b). The recruitment of long-lasting increases in GABA neuron activity contributes to sustained antidepressant effects. Created with BioRender.com.

A major hypothesis for the initial cellular trigger for ketamine’s action it that it first blocks NMDA-R expressed by cortical interneurons, notably, somatostatin (SST) and parvalbumin (PV) GABA neuron subtypes (5, 12). Because these inhibitory neurons exhibit tonic firing, they are thought to be more sensitive to NMDA-R blockade, as tonic activity removes the Mg^2+^ block of the receptor, enabling ketamine to enter the channel pore and block Ca^2+^ entry (12, 40). Inhibition of GABAergic interneuron firing decreases GABA activity, resulting in disinhibition of excitatory pyramidal neurons, and leading to a glutamate burst that drives activity-dependent BDNF release, activation of protein synthesis (via the mTORC1 pathway), new spine formation, and synaptogenesis (2). Supporting this hypothesis, NMDA-R blockade leads to a reduction in the spontaneous firing of putative GABA interneurons in the mPFC, coupled with an enhanced activity of pyramidal neurons at a delayed rate (40). Furthermore, a low concentration of ketamine (1 µM), which approximates brain levels after *in vivo* administration, rapidly decreases inhibitory- and increases excitatory postsynaptic currents (IPSCs and EPSCs, respectively) onto pyramidal neurons in mPFC slices (12).

In this study we measured the dynamics in cellular responses to ketamine using simultaneous dual photometry recordings of GABA neurons and glutamatergic cells in the mPFC. Our findings indicate that ketamine initially produces a decrease in GABAergic neuron activity compared to the vehicle group, and a transient increase in E/I levels (10 min), followed by a long-lasting increase in pyramidal neuron activity (2 h). However, it remains unknown whether the initial decrease in GABAergic transients directly lead to enhanced glutamatergic activity, and more studies addressing cross-correlational dynamics and causality are needed. In a previous study, increased GCaMP6f fluorescence in mPFC pyramidal neurons was observed 30 min after administering a lower dose of ketamine (3 mg/kg), and this study reported more pronounced effects at 30 mg/kg, leading to an immediate increase in pyramidal activity lasting up to 20 min post-injection (41). However, it is noteworthy that the interpretation of that study is limited due to the absence of a vehicle control group. Selective knockdown of a key NMDA-R subunit, GluN2B, in GABA interneurons, but not pyramidal (CaMKII^+^) cells, occluded the behavioral actions of ketamine (12). Optogenetic stimulation of GABA interneurons concurrent with ketamine administration also occluded its behavioral effects (20). Similar cellular mechanisms are observed with other rapid antidepressant candidates, including the muscarinic acetylcholine receptor antagonist scopolamine (14, 29, 30). Stimulating GABA interneuron activity before scopolamine administration blocks its rapid and sustained behavioral effects (30). Likewise, selective knockdown or knockout of muscarinic type 1 (M1) receptors in SST interneurons occludes scopolamine-induced behavioral responses and molecular plasticity at GABAergic and glutamatergic synapses (14, 29). In the same direction, our previous studies have shown that chemogenetic inhibition of either all GAD-positive neurons in the mPFC, or only SST or PV interneuron subtypes, using a higher dose of CNO (2.5 mg/kg, 3x), resulted in fast antidepressant-like responses and molecular changes, mimicking the effects of fast antidepressants (30). Conversely, lower doses of CNO used in the present study (0.5 and 1 mg/kg) are not sufficient to produce behavioral effects on their own, suggesting that a more robust silencing of GAD-positive neurons is required to recapitulate rapid antidepressant actions.

Interestingly, after the initial decrease in GABA activity, we observed a delayed enhancement of GABA function starting 60 min post-ketamine administration, which was further engaged during behavioral tests 24 h and 72 h later, suggesting that additional mechanisms beyond glutamate-mediated plasticity might contribute to ketamine’s actions (Figure 5). Indeed, our chemogenetic experiments provided evidence supporting the involvement of GABAergic interneurons in mediating the effects of ketamine. Notably, while the glutamatergic hypothesis provides a conceptual model for the rapid changes produced by ketamine, it does not account for the reductions in GABA levels and markers observed in cortical areas of human subjects with MDD and chronically stressed animals, which can be restored by rapid antidepressant treatment (10, 15–20, 28, 42–47). Several studies have reported decreases in the expression of SST and proteins related to GABA signaling, such as GAD1—the primary enzyme responsible for synthesizing GABA from glutamate—and multiple subunits of GABA_A_ receptors in cortical tissues obtained from both MDD subjects and stressed animals (48–51). Mice lacking SST (SST-KO) exhibit increased stress-related behaviors, and reduced GAD1 gene expression (52). Importantly, normalization of cortical and plasma GABA levels, as well as GAD1 expression, following antidepressant treatment, is associated with remission of depressive symptoms (21–27).

Consistent with these findings, the antidepressant effects of ketamine are accompanied by robust increase in GABA levels in the mPFC of patients diagnosed with MDD or obsessive-compulsive disorder (19, 53). Ketamine can also restore impaired GABA release in the hippocampus of stressed rats (54). Our previous findings suggest that ketamine and other fast-acting drugs, including scopolamine, increase pre- and postsynaptic markers of GABA signaling in the mPFC 24 h after administration, such as GAD1, the vesicular GABA transporter (VGAT) and/or gephyrin, along with serotonin-induced IPSCs in pyramidal neurons (20). Ketamine can also modulate GABA_A_-R binding in human PFC and increase the activity of extrasynaptic GABA_A_-R in mouse cortex and hippocampus (55, 56). Combined administration of sub-effective doses of muscimol, a selective agonist of GABA_A_ receptors, and ketamine, produces antidepressant-like effects in mice (57). Global deletion or mutation of γ2-GABA-R produces stress-induced phenotypes, which are reversed by a single dose of ketamine or chronic imipramine treatment (58). Ketamine also reverses GABAergic deficits in these animals, increasing inhibitory postsynaptic currents (IPSCs) and gephyrin levels (58). Additionally, disinhibition of somatostatin interneurons by deletion of γ2-GABA-R selectively from these cells results in anxiolytic- and antidepressant-like effects, and confers resilience to stress in male mice, mimicking the effects of ketamine (59, 60). It is worth noting that the experiments in the present study were conducted in single-housed animals that underwent surgeries for cannula placement and/or viral infusion. Social isolation in rodents has been shown to disrupt corticolimbic structures and immune system function, contributing to stress-related behavioral outcomes (61–63). Thus, the single housing may have contributed to baseline stress in these mice that allowed robust changes in response to ketamine administration. However, further experiments using well-validated animal models of stress, such as the CUS, are necessary to evaluate the actions of ketamine compared to a control group (non-stressed).

While the precise mechanism by which ketamine enhances GABA activity remains unclear, one hypothesis is that, following the glutamatergic burst, there is homeostatic self-tuning adaptations to reestablish E/I balance in the mPFC (10). This local reorganization could restore the integrity of signal transfer to target regions by re-establishing correct firing patterns, and thereby promoting antidepressant effects (64) (Figure 5). Glutamate is the primary precursor of GABA as a substrate for GAD, so strengthened inhibition may be a self-limiting mechanism to prevent excitotoxicity in response to excessive excitation. Notably, activation of postsynaptic NMDA-R or stimulation of glutamatergic neurons (CaMKII^+^) engages a positive feedback mechanism culminating in long-term potentiation of dendritic inhibition mediated by SST interneurons in the mPFC (65). Concomitantly, disrupting the glutamate-glutamine cycle depletes GABA neurotransmitter pools (66, 67), restoring balance through decreased inhibition. Another possibility is that ketamine exerts direct effects on the GABAergic system, potentially through its metabolites such as (2R, 6R)-HNK, which has shown ketamine-like antidepressant effects without prompting psychotomimetic responses (38, 68).

Our findings also reveal enhancement of glutamate neuron activity 24 h post-ketamine during the SUST; however, although there was a strong trend towards increased overall glutamatergic activity 72 h later during the NSFT, no effect was found around the bite event, raising the question of whether activity of mPFC CaMKII+ neurons is necessary for the sustained effects of ketamine during specific avoidance-related tasks. In line with our hypothesis, it is possible that activation of glutamatergic neurons is required for the initial cascade of cellular events that culminate in a long-lasting enhancement of dendritic spine number and function (2, 69), while the activity of GABAergic neurons may be involved in fine-tuning these newly formed microcircuits.

Consistent with the involvement of GABA mechanisms in antidepressant responses, new classes of rapid-acting drugs targeting the GABA_A_R as positive or negative allosteric modulators (PAMs and NAMs, respectively) have emerged in recent years. Brexanolone, a positive δ-GABA_A_R modulator and an analogue of the neurosteroid allopregnanolone, has recently been approved for the treatment of postpartum depression (70), marking a significant paradigm shift in the field towards identifying innovative GABAergic compounds for depression therapy. Furthermore, preclinical studies indicate that both PAMs and NAMs of the α5-containing GABA_A_R demonstrate rapid antidepressant-like effects or prevent the behavioral responses induced by chronic stress, comparable to the effects of ketamine (71–74).

Therefore, enhancement of both GABA and glutamate function is likely necessary to restore signal integrity in the mPFC and other limbic regions, culminating in antidepressant states (Figure 5). Our current GABAergic evidence builds upon the findings and conceptual frameworks for antidepressant action described by Duman et al., 2019 (5) and Lusher et al., 2020 (16), providing support for a sequential glutamatergic-GABAergic hypothesis for the actions of ketamine (Figure 5). While the current study provides novel insights into the GABAergic effects of ketamine, we did not evaluate causality between these mechanisms and the enhancement of glutamatergic function. Cross-correlational analyses of GABA and glutamatergic activity, as well as recordings following chemogenetic manipulation of CaMKII^+^ neurons and ketamine treatment, would be valuable additions to investigate the temporal dynamics that could result in disinhibition. Also, further investigations are warranted to pinpoint the recruitment of specific subpopulations of GABA interneurons (e.g., SST, PV) in this biphasic response, as well as potential sex-dependent differences. In conclusion, the current findings suggest that modulating GABAergic function holds promise for developing novel antidepressant medications.

## Author contributions

M.V.F. designed the study, performed the experiments, analyzed the data and wrote the manuscript. F.D. performed experiments and provided technical support. M.R.P. provided scientific input, was involved in data interpretation and analyses, edited and revised the manuscript.

## Acknowledgements

We thank Alexa-Nicole Sliby for helping for her contributions in genotyping and managing the animal colony, and Richard Crouse for his technical assistance with the photometry equipment.

## Data Availability

Individual subject data are shown in the figures, and any unidentified raw data will be provided upon request. MATLAB code is available at https://github.com/katemartian/Photometry_data_processing (34). Any additional information will be provided upon request.

## References

1. Kessler RC, Chiu WT, Demler O, Merikangas KR, Walters EE. Prevalence, severity, and comorbidity of 12-month DSM-IV disorders in the National Comorbidity Survey Replication. Arch Gen Psychiatry. 2005;62(6):617–27.

2. Duman RS, Aghajanian GK, Sanacora G, Krystal JH. Synaptic plasticity and depression: new insights from stress and rapid-acting antidepressants. Nat Med. 2016;22(3):238–49.

3. Trivedi MH, Rush AJ, Wisniewski SR, Nierenberg AA, Warden D, Ritz L, et al. Evaluation of outcomes with citalopram for depression using measurement-based care in STAR*D: implications for clinical practice. Am J Psychiatry. 2006;163(1):28–40.

4. Berman RM, Cappiello A, Anand A, Oren DA, Heninger GR, Charney DS, et al. Antidepressant effects of ketamine in depressed patients. Biol Psychiatry. 2000;47(4):351–4.

5. Duman RS, Sanacora G, Krystal JH. Altered Connectivity in Depression: GABA and Glutamate Neurotransmitter Deficits and Reversal by Novel Treatments. Neuron. 2019;102(1):75–90.

6. Choudary PV, Molnar M, Evans SJ, Tomita H, Li JZ, Vawter MP, et al. Altered cortical glutamatergic and GABAergic signal transmission with glial involvement in depression. Proc Natl Acad Sci U S A. 2005;102(43):15653–8.

7. MacQueen GM, Yucel K, Taylor VH, Macdonald K, Joffe R. Posterior hippocampal volumes are associated with remission rates in patients with major depressive disorder. Biol Psychiatry. 2008;64(10):880–3.

8. Csabai D, Wiborg O, Czeh B. Reduced Synapse and Axon Numbers in the Prefrontal Cortex of Rats Subjected to a Chronic Stress Model for Depression. Front Cell Neurosci. 2018;12:24.

9. Abdallah CG, Jackowski A, Sato JR, Mao X, Kang G, Cheema R, et al. Prefrontal cortical GABA abnormalities are associated with reduced hippocampal volume in major depressive disorder. Eur Neuropsychopharmacol. 2015;25(8):1082–90.

10. Fogaca MV, Duman RS. Cortical GABAergic Dysfunction in Stress and Depression: New Insights for Therapeutic Interventions. Front Cell Neurosci. 2019;13:87.

11. Duman RS, Shinohara R, Fogaca MV, Hare B. Neurobiology of rapid-acting antidepressants: convergent effects on GluA1-synaptic function. Mol Psychiatry. 2019.

12. Gerhard DM, Pothula S, Liu RJ, Wu M, Li XY, Girgenti MJ, et al. GABA interneurons are the cellular trigger for ketamine’s rapid antidepressant actions. J Clin Invest. 2020;130(3):1336–49.

13. Autry AE, Adachi M, Nosyreva E, Na ES, Los MF, Cheng PF, et al. NMDA receptor blockade at rest triggers rapid behavioural antidepressant responses. Nature. 2011;475(7354):91–5.

14. Fogaca MV, Wu M, Li C, Li XY, Duman RS, Picciotto MR. M1 acetylcholine receptors in somatostatin interneurons contribute to GABAergic and glutamatergic plasticity in the mPFC and antidepressant-like responses. Neuropsychopharmacology. 2023;48(9):1277–87.

15. Singh B, Port JD, Voort JLV, Coombes BJ, Geske JR, Lanza IR, et al. A preliminary study of the association of increased anterior cingulate gamma-aminobutyric acid with remission of depression after ketamine administration. Psychiatry Res. 2021;301:113953.

16. Luscher B, Feng M, Jefferson SJ. Antidepressant mechanisms of ketamine: Focus on GABAergic inhibition. Adv Pharmacol. 2020;89:43–78.

17. Czeh B, Vardya I, Varga Z, Febbraro F, Csabai D, Martis LS, et al. Long-Term Stress Disrupts the Structural and Functional Integrity of GABAergic Neuronal Networks in the Medial Prefrontal Cortex of Rats. Front Cell Neurosci. 2018;12:148.

18. Godfrey KEM, Gardner AC, Kwon S, Chea W, Muthukumaraswamy SD. Differences in excitatory and inhibitory neurotransmitter levels between depressed patients and healthy controls: A systematic review and meta-analysis. J Psychiatr Res. 2018;105:33–44.

19. Milak MS, Proper CJ, Mulhern ST, Parter AL, Kegeles LS, Ogden RT, et al. A pilot in vivo proton magnetic resonance spectroscopy study of amino acid neurotransmitter response to ketamine treatment of major depressive disorder. Mol Psychiatry. 2016;21(3):320–7.

20. Ghosal S, Duman CH, Liu RJ, Wu M, Terwilliger R, Girgenti MJ, et al. Ketamine rapidly reverses stress-induced impairments in GABAergic transmission in the prefrontal cortex in male rodents. Neurobiol Dis. 2019:104669.

21. Sanacora G, Mason GF, Rothman DL, Behar KL, Hyder F, Petroff OA, et al. Reduced cortical gamma-aminobutyric acid levels in depressed patients determined by proton magnetic resonance spectroscopy. Arch Gen Psychiatry. 1999;56(11):1043–7.

22. Sanacora G, Gueorguieva R, Epperson CN, Wu YT, Appel M, Rothman DL, et al. Subtype-specific alterations of gamma-aminobutyric acid and glutamate in patients with major depression. Arch Gen Psychiatry. 2004;61(7):705–13.

23. Bhagwagar Z, Wylezinska M, Taylor M, Jezzard P, Matthews PM, Cowen PJ. Increased brain GABA concentrations following acute administration of a selective serotonin reuptake inhibitor. Am J Psychiatry. 2004;161(2):368–70.

24. Goren MZ, Kucukibrahimoglu E, Berkman K, Terzioglu B. Fluoxetine partly exerts its actions through GABA: a neurochemical evidence. Neurochem Res. 2007;32(9):1559–65.

25. Kucukibrahimoglu E, Saygin MZ, Caliskan M, Kaplan OK, Unsal C, Goren MZ. The change in plasma GABA, glutamine and glutamate levels in fluoxetine- or S-citalopram-treated female patients with major depression. Eur J Clin Pharmacol. 2009;65(6):571–7.

26. Karolewicz B, Maciag D, O’Dwyer G, Stockmeier CA, Feyissa AM, Rajkowska G. Reduced level of glutamic acid decarboxylase-67 kDa in the prefrontal cortex in major depression. Int J Neuropsychopharmacol. 2010;13(4):411–20.

27. Dubin MJ, Mao X, Banerjee S, Goodman Z, Lapidus KA, Kang G, et al. Elevated prefrontal cortex GABA in patients with major depressive disorder after TMS treatment measured with proton magnetic resonance spectroscopy. J Psychiatry Neurosci. 2016;41(3):E37–45.

28. Singh B, Port JD, Pazdernik V, Coombes BJ, Vande Voort JL, Frye MA. Racemic ketamine treatment attenuates anterior cingulate cortex GABA deficits among remitters in treatment-resistant depression: A pilot study. Psychiatry Res Neuroimaging. 2022;320:111432.

29. Wohleb ES, Wu M, Gerhard DM, Taylor SR, Picciotto MR, Alreja M, et al. GABA interneurons mediate the rapid antidepressant-like effects of scopolamine. J Clin Invest. 2016;126(7):2482–94.

30. Fogaca MV, Wu M, Li C, Li XY, Picciotto MR, Duman RS. Inhibition of GABA interneurons in the mPFC is sufficient and necessary for rapid antidepressant responses. Mol Psychiatry. 2021;26(7):3277–91.

31. Jendryka M, Palchaudhuri M, Ursu D, van der Veen B, Liss B, Katzel D, et al. Pharmacokinetic and pharmacodynamic actions of clozapine-N-oxide, clozapine, and compound 21 in DREADD-based chemogenetics in mice. Sci Rep. 2019;9(1):4522.

32. Manvich DF, Webster KA, Foster SL, Farrell MS, Ritchie JC, Porter JH, et al. The DREADD agonist clozapine N-oxide (CNO) is reverse-metabolized to clozapine and produces clozapine-like interoceptive stimulus effects in rats and mice. Sci Rep. 2018;8(1):3840.

33. Pothula S, Kato T, Liu RJ, Wu M, Gerhard D, Shinohara R, et al. Cell-type specific modulation of NMDA receptors triggers antidepressant actions. Mol Psychiatry. 2020.

34. Martianova E, Aronson S, Proulx CD. Multi-Fiber Photometry to Record Neural Activity in Freely-Moving Animals. J Vis Exp. 2019(152).

35. Wei C, Han X, Weng D, Feng Q, Qi X, Li J, et al. Response dynamics of midbrain dopamine neurons and serotonin neurons to heroin, nicotine, cocaine, and MDMA. Cell Discov. 2018;4:60.

36. Nathanson JL, Yanagawa Y, Obata K, Callaway EM. Preferential labeling of inhibitory and excitatory cortical neurons by endogenous tropism of adeno-associated virus and lentivirus vectors. Neuroscience. 2009;161(2):441–50.

37. Watakabe A, Ohtsuka M, Kinoshita M, Takaji M, Isa K, Mizukami H, et al. Comparative analyses of adeno-associated viral vector serotypes 1, 2, 5, 8 and 9 in marmoset, mouse and macaque cerebral cortex. Neurosci Res. 2015;93:144–57.

38. Zanos P, Moaddel R, Morris PJ, Georgiou P, Fischell J, Elmer GI, et al. NMDAR inhibition-independent antidepressant actions of ketamine metabolites. Nature. 2016;533(7604):481-6.

39. Tremblay R, Lee S, Rudy B. GABAergic Interneurons in the Neocortex: From Cellular Properties to Circuits. Neuron. 2016;91(2):260–92.

40. Homayoun H, Moghaddam B. NMDA receptor hypofunction produces opposite effects on prefrontal cortex interneurons and pyramidal neurons. J Neurosci. 2007;27(43):11496–500.

41. Hare BD, Pothula S, DiLeone RJ, Duman RS. Ketamine increases vmPFC activity: Effects of (R)- and (S)-stereoisomers and (2R,6R)-hydroxynorketamine metabolite. Neuropharmacology. 2020;166:107947.

42. Prescot A, Sheth C, Legarreta M, Renshaw PF, McGlade E, Yurgelun-Todd D. Altered Cortical GABA in Female Veterans with Suicidal Behavior: Sex Differences and Clinical Correlates. Chronic Stress (Thousand Oaks). 2018;2.

43. Kanes SJ, Colquhoun H, Doherty J, Raines S, Hoffmann E, Rubinow DR, et al. Open-label, proof-of-concept study of brexanolone in the treatment of severe postpartum depression. Hum Psychopharmacol. 2017;32(2).

44. Chowdhury GM, Zhang J, Thomas M, Banasr M, Ma X, Pittman B, et al. Transiently increased glutamate cycling in rat PFC is associated with rapid onset of antidepressant-like effects. Mol Psychiatry. 2017;22(1):120–6.

45. Perrine SA, Ghoddoussi F, Michaels MS, Sheikh IS, McKelvey G, Galloway MP. Ketamine reverses stress-induced depression-like behavior and increased GABA levels in the anterior cingulate: an 11.7 T 1H-MRS study in rats. Prog Neuropsychopharmacol Biol Psychiatry. 2014;51:9–15.

46. Guilloux JP, Douillard-Guilloux G, Kota R, Wang X, Gardier AM, Martinowich K, et al. Molecular evidence for BDNF- and GABA-related dysfunctions in the amygdala of female subjects with major depression. Mol Psychiatry. 2012;17(11):1130–42.

47. Sibille E, Morris HM, Kota RS, Lewis DA. GABA-related transcripts in the dorsolateral prefrontal cortex in mood disorders. Int J Neuropsychopharmacol. 2011;14(6):721–34.

48. Merali Z, Du L, Hrdina P, Palkovits M, Faludi G, Poulter MO, et al. Dysregulation in the suicide brain: mRNA expression of corticotropin-releasing hormone receptors and GABA(A) receptor subunits in frontal cortical brain region. J Neurosci. 2004;24(6):1478–85.

49. Sequeira A, Klempan T, Canetti L, ffrench-Mullen J, Benkelfat C, Rouleau GA, et al. Patterns of gene expression in the limbic system of suicides with and without major depression. Mol Psychiatry. 2007;12(7):640–55.

50. Klempan TA, Sequeira A, Canetti L, Lalovic A, Ernst C, ffrench-Mullen J, et al. Altered expression of genes involved in ATP biosynthesis and GABAergic neurotransmission in the ventral prefrontal cortex of suicides with and without major depression. Mol Psychiatry. 2009;14(2):175–89.

51. Luscher B, Shen Q, Sahir N. The GABAergic deficit hypothesis of major depressive disorder. Mol Psychiatry. 2011;16(4):383–406.

52. Lin LC, Sibille E. Somatostatin, neuronal vulnerability and behavioral emotionality. Mol Psychiatry. 2015;20(3):377–87.

53. Rodriguez CI, Kegeles LS, Levinson A, Ogden RT, Mao X, Milak MS, et al. In vivo effects of ketamine on glutamate-glutamine and gamma-aminobutyric acid in obsessive-compulsive disorder: Proof of concept. Psychiatry Res. 2015;233(2):141–7.

54. Tornese P, Sala N, Bonini D, Bonifacino T, La Via L, Milanese M, et al. Chronic mild stress induces anhedonic behavior and changes in glutamate release, BDNF trafficking and dendrite morphology only in stress vulnerable rats. The rapid restorative action of ketamine. Neurobiol Stress. 2019;10:100160.

55. Heinzel A, Steinke R, Poeppel TD, Grosser O, Bogerts B, Otto H, et al. S-ketamine and GABA-A-receptor interaction in humans: an exploratory study with I-123-iomazenil SPECT. Hum Psychopharmacol. 2008;23(7):549–54.

56. Wang DS, Penna A, Orser BA. Ketamine Increases the Function of gamma-Aminobutyric Acid Type A Receptors in Hippocampal and Cortical Neurons. Anesthesiology. 2017;126(4):666–77.

57. Rosa PB, Neis VB, Ribeiro CM, Moretti M, Rodrigues AL. Antidepressant-like effects of ascorbic acid and ketamine involve modulation of GABAA and GABAB receptors. Pharmacol Rep. 2016;68(5):996–1001.

58. Ren Z, Pribiag H, Jefferson SJ, Shorey M, Fuchs T, Stellwagen D, et al. Bidirectional Homeostatic Regulation of a Depression-Related Brain State by Gamma-Aminobutyric Acidergic Deficits and Ketamine Treatment. Biol Psychiatry. 2016;80(6):457–68.

59. Jefferson SJ, Feng M, Chon U, Guo Y, Kim Y, Luscher B. Disinhibition of somatostatin interneurons confers resilience to stress in male but not female mice. Neurobiol Stress. 2020;13:100238.

60. Fuchs T, Jefferson SJ, Hooper A, Yee PH, Maguire J, Luscher B. Disinhibition of somatostatin-positive GABAergic interneurons results in an anxiolytic and antidepressant-like brain state. Mol Psychiatry. 2017;22(6):920–30.

61. Du Preez A, Law T, Onorato D, Lim YM, Eiben P, Musaelyan K, et al. The type of stress matters: repeated injection and permanent social isolation stress in male mice have a differential effect on anxiety- and depressive-like behaviours, and associated biological alterations. Transl Psychiatry. 2020;10(1):325.

62. Takatsu-Coleman AL, Patti CL, Zanin KA, Zager A, Carvalho RC, Borcoi AR, et al. Short-term social isolation induces depressive-like behaviour and reinstates the retrieval of an aversive task: mood-congruent memory in male mice? J Psychiatry Neurosci. 2013;38(4):259–68.

63. Agis-Balboa RC, Pinna G, Pibiri F, Kadriu B, Costa E, Guidotti A. Down-regulation of neurosteroid biosynthesis in corticolimbic circuits mediates social isolation-induced behavior in mice. Proc Natl Acad Sci U S A. 2007;104(47):18736–41.

64. Turrigiano GG, Nelson SB. Homeostatic plasticity in the developing nervous system. Nat Rev Neurosci. 2004;5(2):97–107.

65. Chiu CQ, Martenson JS, Yamazaki M, Natsume R, Sakimura K, Tomita S, et al. Input-Specific NMDAR-Dependent Potentiation of Dendritic GABAergic Inhibition. Neuron. 2018;97(2):368–77 e3.

66. Liang SL, Carlson GC, Coulter DA. Dynamic regulation of synaptic GABA release by the glutamate-glutamine cycle in hippocampal area CA1. J Neurosci. 2006;26(33):8537–48.

67. Rae C, Hare N, Bubb WA, McEwan SR, Broer A, McQuillan JA, et al. Inhibition of glutamine transport depletes glutamate and GABA neurotransmitter pools: further evidence for metabolic compartmentation. J Neurochem. 2003;85(2):503–14.

68. Fukumoto K, Fogaca MV, Liu RJ, Duman C, Kato T, Li XY, et al. Activity-dependent brain-derived neurotrophic factor signaling is required for the antidepressant actions of (2R,6R)-hydroxynorketamine. Proc Natl Acad Sci U S A. 2019;116(1):297–302.

69. Li N, Lee B, Liu RJ, Banasr M, Dwyer JM, Iwata M, et al. mTOR-dependent synapse formation underlies the rapid antidepressant effects of NMDA antagonists. Science. 2010;329(5994):959-64.

70. Morrow AL, Balan I, Boero G. Mechanisms Underlying Recovery From Postpartum Depression Following Brexanolone Therapy. Biol Psychiatry. 2022;91(3):252–3.

71. Troppoli TA, Zanos P, Georgiou P, Gould TD, Rudolph U, Thompson SM. Negative Allosteric Modulation of Gamma-Aminobutyric Acid A Receptors at alpha5 Subunit-Containing Benzodiazepine Sites Reverses Stress-Induced Anhedonia and Weakened Synaptic Function in Mice. Biol Psychiatry. 2022;92(3):216–26.

72. Xiong Z, Zhang K, Ishima T, Ren Q, Chang L, Chen J, et al. Comparison of rapid and long-lasting antidepressant effects of negative modulators of alpha5-containing GABAA receptors and (R)ketamine in a chronic social defeat stress model. Pharmacol Biochem Behav. 2018;175:139–45.

73. Piantadosi SC, French BJ, Poe MM, Timic T, Markovic BD, Pabba M, et al. Sex-Dependent Anti-Stress Effect of an alpha5 Subunit Containing GABAA Receptor Positive Allosteric Modulator. Front Pharmacol. 2016;7:446.

74. Zanos P, Nelson ME, Highland JN, Krimmel SR, Georgiou P, Gould TD, et al. A Negative Allosteric Modulator for alpha5 Subunit-Containing GABA Receptors Exerts a Rapid and Persistent Antidepressant-Like Action without the Side Effects of the NMDA Receptor Antagonist Ketamine in Mice. eNeuro. 2017;4(1).

