## Supplementary material for "Effects of ketamine on GABAergic and glutamatergic activity in the mPFC: biphasic recruitment of GABA function in antidepressant-like responses"

**Supplementary Figures**

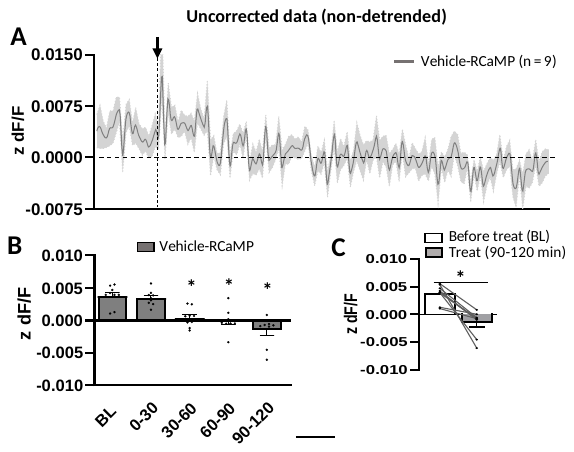

**Supplementary Figure 1 – Uncorrected (non-detrended) photometry data from mPFC GABAergic neurons immediately after vehicle treatment.** (A-B) There was a gradual photobleaching of RCaMP fluorescence over the 2-h recording period that was not corrected using the 405 isobestic signal (Repeated measures ANOVA, F_time 4,32_ = 15.33, p ≤ 0.05). (C)
Individual before-and-after data plots demonstrate a significant decrease in calcium transients at the end of the recording compared to the period before injection (baseline, BL) (paired Student’s t-test, t_8_ = 5.11, p ≤ 0.05). Graphs are presented as mean ± standard error.

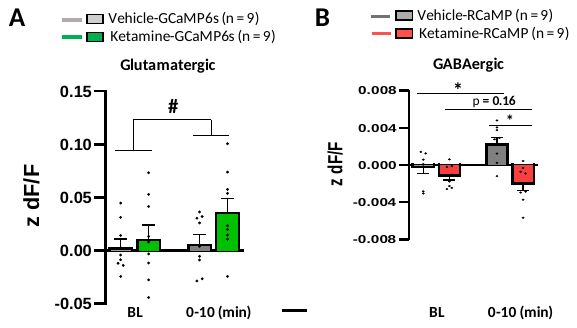

**Supplementary Figure 2 – Photometry transients recorded from mPFC CaMKII^+^ and Gad^+^ neurons during 10 min following treatments.** (A) Main effect of time on the activity of mPFC CaMKII+ neurons in mice treated with either vehicle or ketamine (F_time 1,8_ = 7.30, p < 0.05). (B) Effect of ketamine on photometry transients from mPFC GABAergic neurons (F_treat 1,8_ = 33.58, p < 0.05; F_time 1,8_ = 5.61, p < 0.05; F_interaction 1,8_ = 15.35; p <0.05). # p < 0.05, main effect of time, Repeated measures ANOVA; *p < 0.05, Repeated measures ANOVA followed by Sidak.

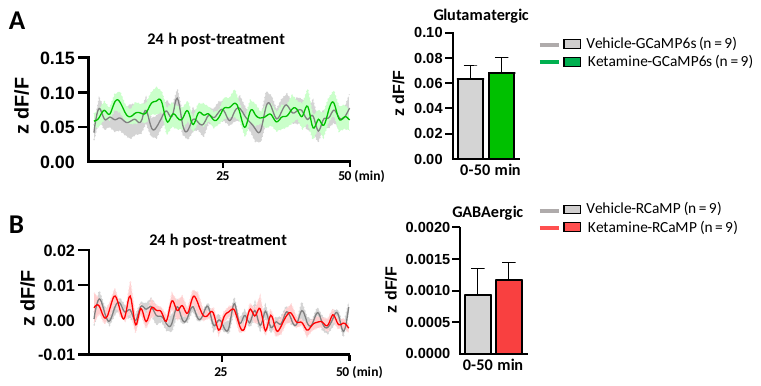

**Supplementary Figure 3. Photometry recordings from glutamate and GABA neurons 24 h after ketamine treatment.** Animals were treated with either vehicle or ketamine and on the next day underwent recordings for 50 min before the behavioral test. (A) Ketamine did not produce baseline changes in glutamatergic (t_16_ = 0.30, p > 0.05) or (B) GABAergic (t_16_ = 0.49, p > 0.05) calcium transients 24 h following administration. Photometry results are computed as average z-scored dF/F, and graphs are presented as mean ± standard error. Student’s t-test (two-tailed), n = 9/group.

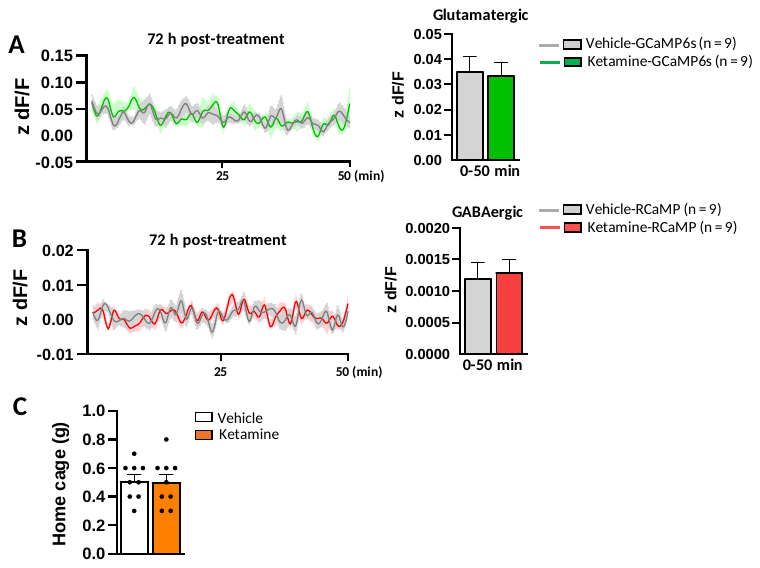

**Supplementary Figure 4. Photometry recordings from glutamate and GABA neurons 72 h after ketamine treatment.** Animals were treated with either vehicle or ketamine and, 72 h later, underwent recordings for 50 min before the behavioral test. (A) Ketamine did not produce baseline changes in glutamatergic (t_16_ = 0.18, p > 0.05) or (B) GABAergic (t_16_ = 0.29, p > 0.05) calcium transients 72 h following administration. (C) Ketamine did not change home cage food consumption in the novelty suppressed feeding test (t_16_ = 0.16, p > 0.05). Photometry results are computed as average z-scored dF/F, and graphs are presented as mean ± standard error. Student’s t-test (two-tailed), n = 9/group.

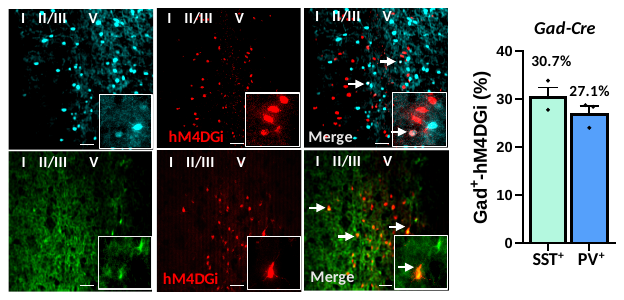

**Supplementary Figure 5. Quantification of co-localized Gad-hM4DGi expression and SST or PV cells.** Gad-cre mice received bilateral infusion of a Cre-dependent hM4DGi virus into the mPFC for quantification of co-labeling with PV or SST cells (n = 3 mice). Magnification: 20X; inset: 40X. Graphs are presented as mean ± standard error.

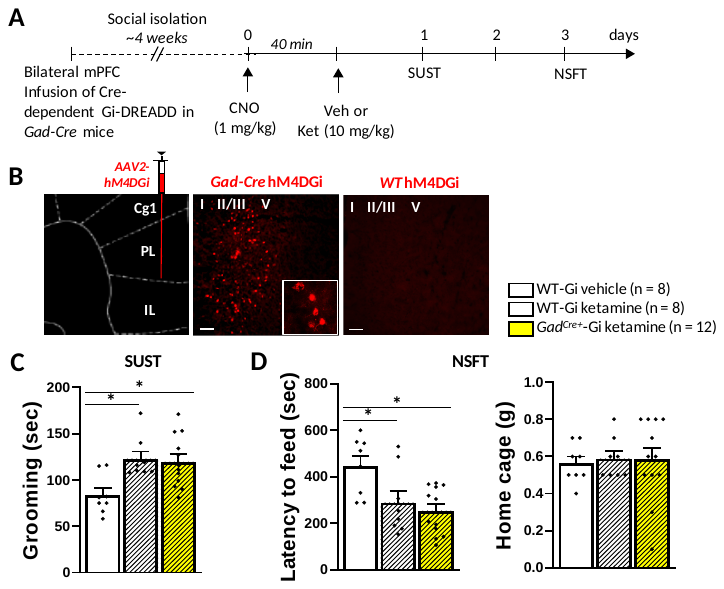

**Supplementary Figure 6.** **Chemogenetic inhibition of Gad^+^ neurons before ketamine treatment.** (A) Timeline for surgery, treatments, and behavioral testing. (B) Representative images of the mPFC from *Gad1-Cre* and WT (Cre-negative littermate) mice that received bilateral infusions of hM4DGi virus (magnification: 10X; inset: 40X). (C) CNO (1 mg/kg) administration 40 min before ketamine did not impact its behavioral effects the sucrose splash test (SUST, F_2,25_ = 6.47) and (D) novelty supressed feeding test (NSFT, F_2,25_ = 6.20), 24 and 72 hours following treatment, respectively. There was no change in home cage food comsumption (F_2,25_ = 0.05). Graphs are presented as mean ± standard error. * p < 0.05, different from the vehicle-treated group. One-way ANOVA followed by Duncan test.
